# Community and leaf-level controls of carbon fluxes in an invaded NZ alpine tussock grassland

**DOI:** 10.64898/2026.04.21.719903

**Authors:** Indira V. Leon-Garcia, Aimée T. Classen, Julie R. Deslippe

## Abstract

Climate warming and plant invasion are reshaping alpine communities, yet their combined effects on ecosystem carbon (C) balance remain poorly understood. We studied how warming and invasion interact to influence community level C fluxes, and the mechanisms underpinning these responses. Using field experiments manipulating warming and invader removal in two New Zealand alpine grasslands dominated by functionally distinct invaders (a shrub, *Calluna vulgaris*, and a forb, hawkweeds), we quantified net ecosystem exchange (NEE), gross primary productivity (GPP) and ecosystem respiration (ER), alongside community, trait and abiotic factors. Warming did not enhance C uptake at either site, and we found no evidence of synergistic warming × invasion effects. Instead, responses were invader-dependent. At the shrub-invaded site, warming increased net C loss through elevated respiration without corresponding gains in GPP. In contrast, fluxes at the forb-invaded site were largely insensitive to treatments. Variation in C fluxes were driven by abiotic conditions and trait dominance at the shrub site, whereas species diversity (richness and evenness) exerted stronger control at the forb site. Our findings demonstrate that the impacts of invasion on ecosystem C balance depend on invader functional identity, with contrasting roles for trait dominance and biodiversity in regulating ecosystem function under warming.

## Introduction

Climate change is a major driver of biodiversity change in alpine ecosystems (Sala et al., 2000). High mountain ecosystems are typically characterized by low or patchy plant cover and low productivity (Chen et al., 2017; Parmesan, 2006), largely due to the physiological constraints imposed by cold temperatures that limit carbon uptake and turnover (Llambí et al., 2018; Stuart Chapin III et al., 2009). However, alpine ecosystems are warming rapidly (Chelli et al., 2017; Jabis, 2018), and increasing anthropogenic disturbance is likely to amplify invasion pressure as environmental filters are relaxed (Carboni et al., 2018; Kalwij et al., 2008). Because colder ecosystems may respond more strongly to warming (Wang et al., 2012; Zhang et al., 2015) and store substantial carbon belowground (Buytaert et al., 2011; Curiel Yuste et al., 2017), it is critical to understand how carbon cycling will change as community composition shifts.

Carbon exchange between plants and the atmosphere is mediated by photosynthesis and respiration, with their balance reflected in net ecosystem exchange (NEE), where negative values indicate net carbon uptake and positive values indicate carbon loss to the atmosphere (Schulze et al., 2021; Wood, 2021). Rising atmospheric CO_2_ may enhance photosynthesis and increase carbon uptake, but only where nutrient limitation does not constrain growth (Chen et al., 2017; Falkowski et al., 2000). Importantly, species- and functional groups differ in their responses to warming and elevated CO_2_. For example, warming effects on leaf-level carbon fluxes depend on leaf nitrogen content in grasses, but not in forbs (Chen et al., 2017; Premke et al., 2016; Zhou et al., 2021). These differences highlight the importance of linking canopy-scale fluxes to species traits and physiology to improve predictions of ecosystem carbon dynamics under climate change (Becklin et al., 2021; Friend et al., 2007; Grace, 2004).

Plant functional diversity plays an important role in regulating ecosystem C balance because species vary in their ability to capture, store and release carbon (Conti & Diaz, 2013). Two components of biodiversity are commonly used to explain variation in C fluxes: vegetation quantity (e.g., biomass or abundance; Lohbeck et al., 2015) and vegetation quality (trait dominance and functional diversity; Conti & Diaz, 2013; Diaz et al., 2007; Prado-Junior et al., 2016). These correspond to distinct but not mutually exclusive hypotheses. The biomass ratio hypothesis (Grime, 1998) posits that ecosystem function is driven by the traits of dominant species; such that community weighted mean (CWM) traits strongly predict carbon fluxes. In contrast, the niche complementary hypothesis (Trenbath, 1974), suggests that greater species or trait diversity enhances resource-use efficiency and productivity (Díaz et al., 2011; Tilman, 1999; Zhang et al., 2012). These mechanisms align with the selection and complementarity effects described by Loreau in 2010 (Loreau, 2010; Loreau & Hector, 2001), and can operate independently or interact depending on environmental context and community composition (Mouillot et al., 2011; Shen et al., 2016). In grasslands, aboveground biomass and trait dominance are often strong predictors of carbon fluxes (Everwand et al., 2014). However, climate-driven shifts in species composition and trait diversity can also alter C balance. For example, warming increased NEE by 31% on the Tibetan plateau through shifts towards graminoids and legumes, which increased soil nitrogen availability (Chen et al 2017). Such responses illustrate how changes in biomass and functional traits scale to ecosystem-level C dynamics under climate change (Chen et al., 2017).

Biological invasions can further modify carbon cycling by altering both community composition and resource availability. Invaders often increase total biomass and productivity (Vought et al., 2026), while reducing native diversity (Castro-Díez et al., 2016; Gibbons et al., 2017; Poland et al., 2021). These effects arise because many invasive species exhibit high productivity, phenotypic plasticity, and resource-use efficiency (Van Kleunen et al., 2010), with traits such as clonality, height, and extended phenology associated with invasiveness (Dassonville et al., 2008; Goodell & Parker, 2017; Milanović et al., 2020). As invaders become dominant, they can increase decomposition rates (Liao et al., 2008), and alter soil biota (Moyle & Deslippe, 2024), while reducing resource availability for native species (Castro-Díez et al., 2019; Helsen et al., 2018; Peltzer et al., 2009; Vilà et al., 2011). This dominance-driven increase in ecosystem function under reduced diversity (Flombaum et al., 2017; Livingstone et al., 2020) highlights a key pathway through which invasion influences carbon fluxes. However, invasion impacts are context-dependent, varying with invader traits, community composition, and environmental conditions (Gaertner et al., 2009; Novoa et al., 2020; Sheppard & Lüpke, 2024). Functionally distinct invaders may exert particularly strong effects when they replace key functional groups (Allison & Vitousek, 2004; Vitousek, 1986), underscoring the need to compare invader types when predicting ecosystem responses.

In Aotearoa New Zealand (hereafter ‘Aotearoa’), alpine ecosystems are experiencing rapid warming and increasing invasion pressure. Warming and drying threaten native alpine flora, with 40-70% of species at risk of extinction (Halloy & Mark, 2003). Concurrently, invasive species such as the European shrub *Calluna vulgaris* are transforming native plant community composition (Chapman & Bannister, 1990; Moyle & Deslippe, 2024). In its native range, *C. vulgaris* dominates upland heaths and sequesters more carbon than nearby grasslands, partly due to lower metabolic rates and reduced litter turnover (Quin et al., 2015). In Aotearoa, it exhibits high phenological plasticity under warming, enhancing its competitive advantage (Giejsztowt et al., 2020), and suppresses native species by disrupting pollination and mycorrhizal associations (Giejsztowt et al., 2020; Moyle & Deslippe, 2024). Rosetted hawkweeds (*Pilosella officinarum* and *Hieracium radicata*) represent a second invader type expanding in alpine zones. Hawkweeds displace native short-tussock grasslands (Steer & Norton, 2013), potentially altering ecosystem function through the loss of functionally distinct species.

To test how warming and invasion independently and interactively affect ecosystem function, we conducted a factorial field experiment manipulating warming and invader removal at two alpine sites in Aotearoa: one dominated by *C. vulgaris* and the other by rosetted hawkweeds. Our 2×2 design included ambient control (AC), warming control (WC), ambient removal (AR), and warming and removal (WR) treatments. We quantified plant community composition, functional traits, and ecosystem carbon fluxes (NEE, gross primary productivity (GPP), and ecosystem respiration (ER)). We tested two hypotheses. First (Hyp1), we predicted that warming would increase carbon uptake (more negative NEE and GPP) by enhancing plant productivity, but that removal effects would differ by site. At the hawkweeds site, we expected removal (AR, WR) to increase carbon uptake by reducing competition and increasing native diversity. At the heather site, we expected removal would reduce carbon uptake by eliminating a dominant, high-biomass shrub and increase respiration through root decomposition, leading to net carbon loss. Second (Hyp2), we tested which components of biodiversity best predicted carbon fluxes. We expected vegetation quantity (biomass, percent cover, trait dominance) to dominate carbon flux at the heather site (biomass ratio hypothesis), whereas functional and species diversity would better predict carbon fluxes at the hawkweed site (niche complementarity hypothesis).

## Methods

### Study sites

The study took place in two alpine tussock-grassland communities within or nearby Tongariro National Park (TNP) Aotearoa. The first site (“heather site”) is located on the lee slopes of Mount Ruapehu (39.19 S, 175.59 E, 1070 m.a.s.l.) and is dominated by the invasive dwarf shrub *Calluna vulgaris*. The second site (“hawkweed site”) is located on the southern slopes of Mount Ruapehu (39.38 S, 175.69 E at 1340 m a.s.l.) and is invaded by rosetted forbs (*Pilosella officinarum* and *Hypochaeris radicata*, hereafter “hawkweeds”). Both sites are characterised by native tussock grasslands with co-occurring ericaceous shrubs (e.g. *Dracophyllum sp*.) and native grasses (e.g. *Chionochloa* rubra., *Poa spp*.). Hawkweeds occur uncommonly and in low abundance at the heather site, while heather does not occur at the hawkweed site. Additional details on the two study sites are described in Leon-Garcia & Deslippe, 2025.

At each site, we implemented a factorial field experiment manipulating warming and invader removal. The core design consisted of a 2×2 factorial combination of warming (ambient vs. warmed) and invader removal (present vs. removed), yielding four treatments: ambient control (AC), warming control (WC), ambient removal (AR), and warming plus removal (WR, n=8 per treatment). Warming was applied with hexagonal open top chambers (OTCs; ITEX design) constructed from plexiglass, which increased mean air temperature by approximately +1.7 °C (Giejsztowt et al. 2020). To better isolate treatment effects on carbon flux, we included two additional ambient treatments (n = 6): (1) uninvaded control plots to account for invasion effects (A), and (2) plots where a native species of similar growth form to the invader was removed to control for removal effects (ANR). At the heather site the ANR treatment targeted *Dracophyllum subulatum*, while at the hawkweed site it targeted *Gentianella bellidifolia*. Removal treatments were implemented by clipping aboveground biomass of target species to ground level and were repeated as needed throughout the growing season (typically 2–3 times per year). In total, each site comprised 44 plots (Fig 2). At the time of carbon flux measurements, the heather experiment had been running for eight years and the hawkweed experiment for two years. Air temperature, soil moisture, and soil temperature were recorded continuously using TMS-4 dataloggers (TOMST, Czech Republic) in a subset of plots at each site.

**Figure 1.**
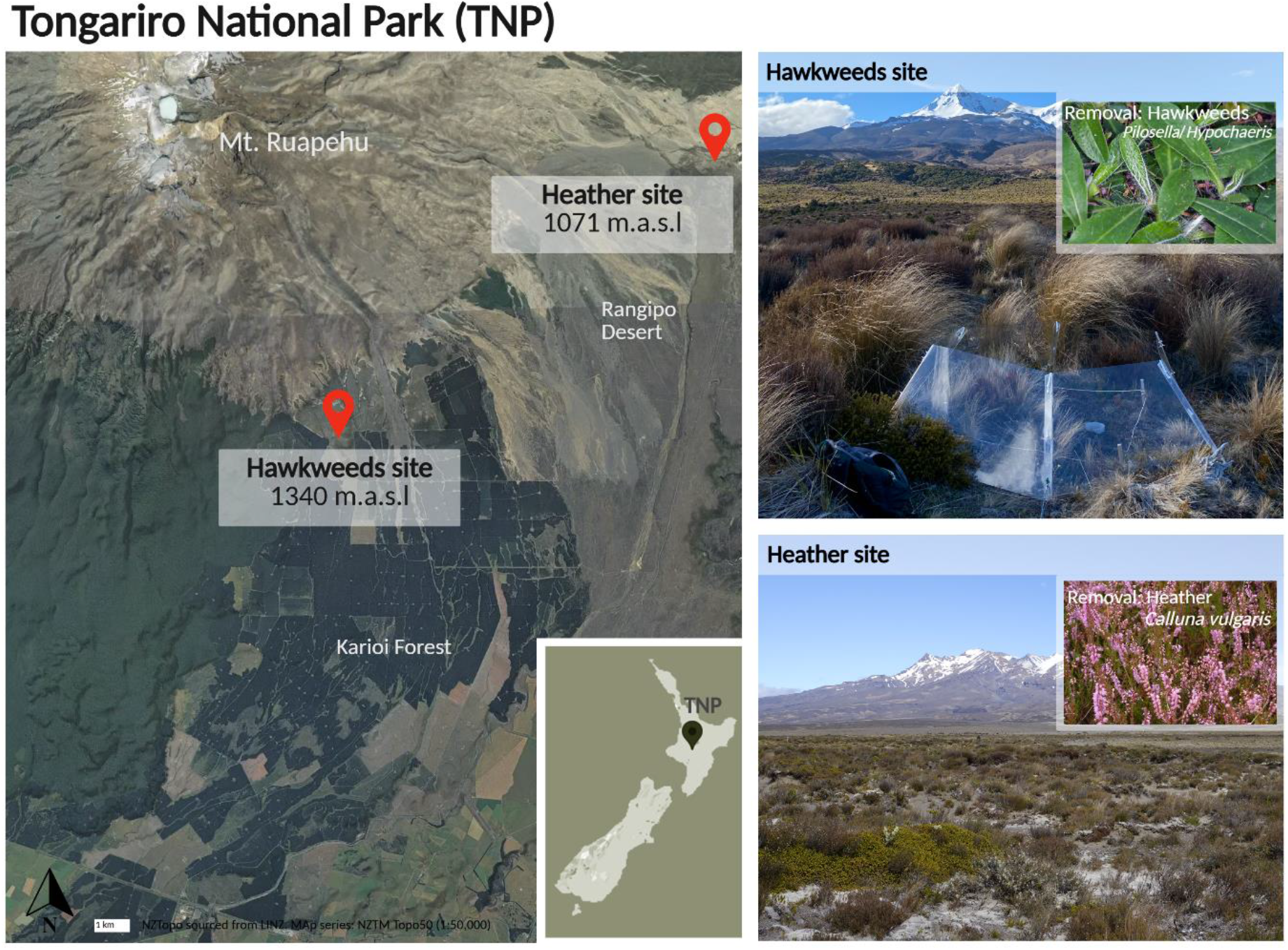
Locations of the two study sites on Mount Ruapehu in Tongariro National Park (TNP), Aotearoa. Different invaders are present at each site: the European ericaceous shrub, heather (*Calluna vulgaris*), invades the eastern leeward slopes (bottom right) while hawkweed (*Pilosella officinarum* and *Hypochaeris radicata*) forbs invade the Southern site (top right).

**Figure 2.**
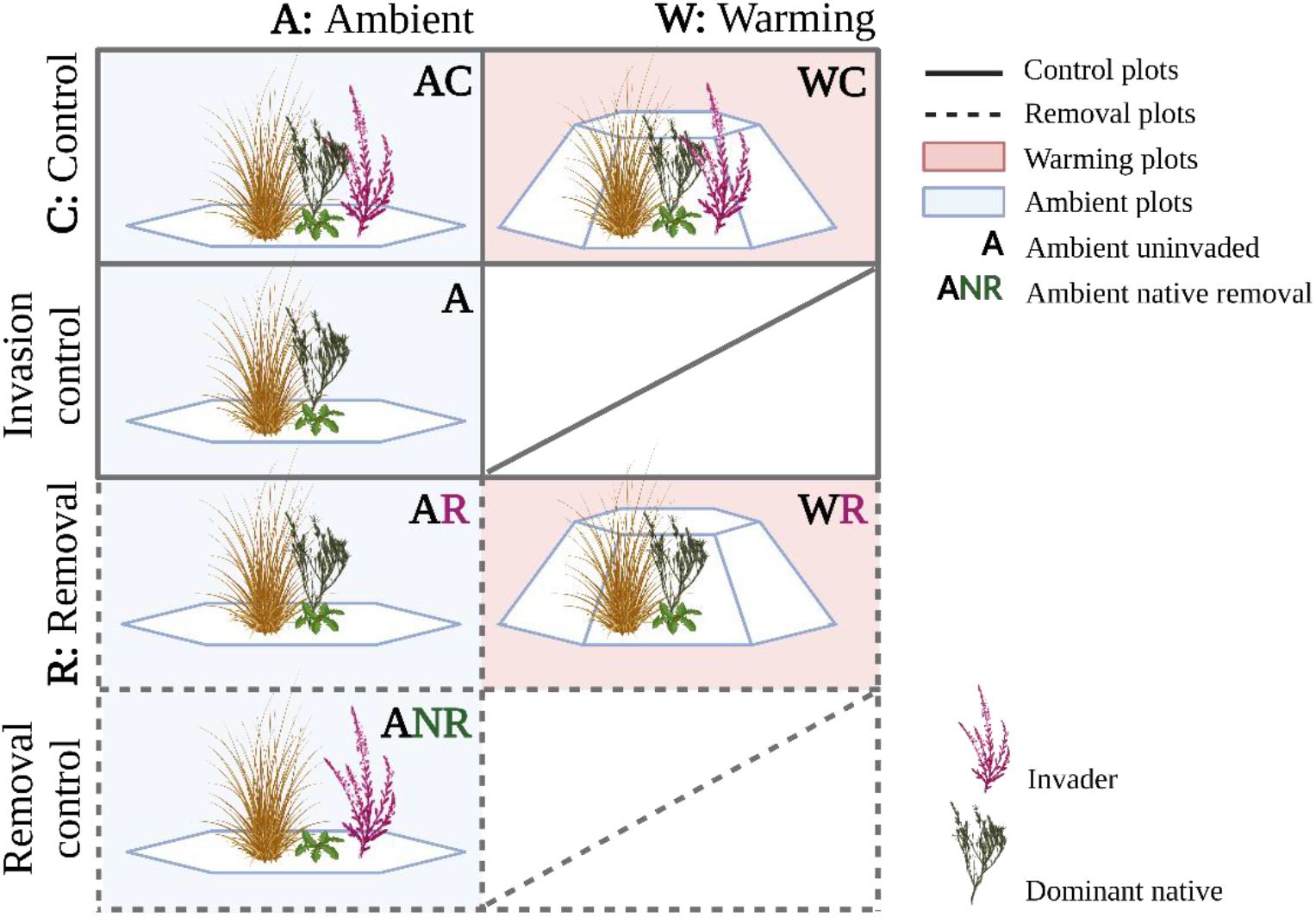
Experimental design at heather and hawkweed sites. Warming is applied with open top chambers (OTC) and invader removal is applied by clipping plants to ground level. The ANR treatments targeted the ericaceous shrub *Dracophyllum subulatum* at the heather site and the forb *Gentianella bellidifolia* at the hawkweed site. The heather site was established in 2015 while the hawkweed site was established in 2022.

### Ecosystem carbon flux measurements

To assess the effects of warming and invasive species on ecosystem carbon fluxes we measured net ecosystem exchange (NEE), gross primary productivity (GPP), and ecosystem respiration (ER) at peak season biomass in January and February 2024. We measured the change in carbon dioxide concentration at three light intensities and in the dark (ER) inside an airtight chamber placed over the top of the vegetation and sealed to the ground. CO_2_ concentrations were recorded using an infrared gas analyzer (LI-COR 7500, Lincoln, Nebraska, USA), and the components of carbon flux were calculated as described by Vought et al., 2026. Full details of the carbon flux measurements and calculation also appear in supplementary material.

### Plant composition, diversity and focal species selection

To evaluate how ecosystem C fluxes were influenced by plant composition under treatments we assessed plant cover and species and functional diversities in all plots. Plant community composition was assessed at peak season (January–March 2024) at both sites using visual estimates of percent cover within 1 m^2^ quadrats as described by Leon-Garcia et al., 2025. All species were recorded, along with bare soil and litter. Total plant cover (excluding soil and litter) and relative species cover were calculated per plot. We also calculated species richness (S), Shannon diversity (H), and evenness (J) for each plot.

Species contributing ≥80% of total cover and occurring consistently across treatments were targeted as focal species for trait measurements. At the heather site, focal species included the invader *Calluna vulgaris* and three dominant natives (*Dracophyllum subulatum, Chionochloa rubra, Veronica tetragona*). At the hawkweed site, invasive forbs (*Pilosella officinarum, Hypochaeris radicata*) were grouped as “hawkweeds”, alongside four dominant natives (*Dracophyllum subulatum, Poa cita, Wahlenbergia pygmaea, Gentianella bellidifolia*).

### Focal species functional trait measurements

We measured leaf traits associated with carbon uptake and water use: leaf thickness (L_th_), relative water content (RWC), specific leaf area (SLA), stomatal density (SD), leaf δ^13^C isotope, maximum light-saturated photosynthetic rate (A_max_) and leaf dark respiration (R_D_). Traits were measured on fully expanded leaves following Pérez-Harguindeguy et al., 2013. Fresh leaves were used for RWC, L_*th*_, and SD measurements. RWC was calculated from fresh, saturated, and dry mass. Leaf thickness was measured with a microtome (excluding midribs), and stomatal density from nail polish impressions. Gas exchange (A_max_ and R_D_) was measured in the field using a LI-COR 6400XT (LI-COR, Lincoln, NE, USA) between 08:00 and 13:00 at 1500 µmol m^−2^ s^−1^PAR. Fluxes were corrected for leaf area differences among species. For traits expressed as negative values (δ^13^C, R_D_), absolute values were used to ease interpretation. Normalized Difference Vegetation Index (NDVI) was measured using a handheld GreenSeeker sensor. Twelve readings per plot were averaged at peak productivity (January–February). NDVI correlates strongly with GPP at these sites (Sundqvist et al., 2020).

Trait data were used to calculate community weighted means (CWM) and functional diversity (Rao’s quadratic entropy, FD_Q_) at the plot level, weighted by species’ relative cover. CWMs were calculated following Garnier et al. 2007:

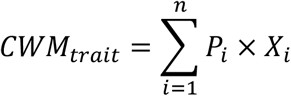

were *P*_*i*_ is species relative abundance and *X*_*i*_ the trait values (including the invader). FD_Q_ incorporated pairwise trait distances among species and their abundances. FD_Q_ was calculated following Botta-Dukát, 2005:

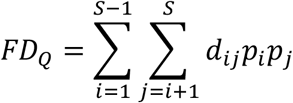

were *d*_*ij*_ is the distance between two species in the community and *p* their relative abundance calculated from the percent cover data. The metric captures multi-trait functional differentiation and reflects shifts in trait diversity associated with invasion. Calculations were conducted in R using standard ‘tidyverse’ workflows and the SYNCSA package.

### Data analysis

To test our first hypothesis on treatment effects on carbon fluxes (NEE, GPP, ER), we first reduced predictor redundancy using principal components analysis (PCA). Predictors included: biomass-related variables (dominant and invader cover, estimated biomass, and NDVI), community trait metrics (CWM and FD_Q_), species diversity indices (richness, Shannon’s diversity, evenness), and abiotic variables (mean and maximum air temperature, soil moisture). We retained principal components explaining most variance and selected the ten highest-loading predictors, ensuring cumulative variance exceeded 80% by PC2 (Jackson, 1991). For this analysis we employed the *FactoMineR, factoextra* and *ggcorrplot* packages in R. PC1 and PC2 were subsequently used to summarise multivariate variation and, where appropriate, included as predictors.

Treatment effects were assessed using linear mixed-effects models (LMMs) for each flux (NEE, GPP, ER), following Everwand et al., 2014, and Diaz et al., 2007. Predictors were added hierarchically: abiotic terms; species composition terms; community traits, and models were. simplified using Akaike Information Criterion corrected (AICc) and likelihood ratio tests to obtain the most parsimonious structure (Diaz et al., 2007). Warming and removal were included as fixed effects, with plots nested within blocks as random effects:

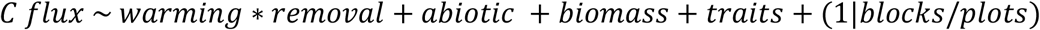

Abiotic predictors were air temperature or soil moisture; biomass terms included NDVI, total cover, and above-ground biomass; trait predictors comprised CWM and FD_Q_. PCA axes were included when they improved model fit. For parametrization, we implemented logarithmic transformations for the response C fluxes as needed. Response variables were log-transformed where necessary. Models were fitted with the ‘lmer’ function on the *lme4* R package with model selection via likelihood ratio tests using the ‘lrtest’ function in *lmtest*. To explore invader effects on native community composition and diversity, we we restricted analyses to plots where invaders were present (AC, WC). We fitted LMMs and a correlation matrix using the same predictor set. Correlations were calculated using the ‘correlate’ function in the *corrplot* package, with significance assessed using ‘cor.mtest’ and visualised via MDS clustering.

To test our second hypothesis on the drivers of C fluxes, we used structural equation modelling (SEM) with path analysis for each flux (NEE, GPP, ER) to quantify the magnitude and direction of the direct and indirect effects, and to identify relationships among predictors. We based our *a priori* path models on a combination of the trait-based productivity (TBP) theory proposed by Yan et al., 2023, and models employed by other recent similar studies integrating the effect of biodiversity on ecosystem functioning (De Long et al., 2019; Loreau, 2010; Milcu et al., 2014). The conceptual model evaluates how abiotic factors (resource availability) and plant diversity influence carbon fluxes directly and indirectly via plant functional traits (Fig. 3). Specifically, we expected invasive species biomass and traits to modify community composition and resource uptake, thereby affecting carbon uptake. Predictors included variables identified as influential in the PCA and used in LMMs, with PC1 and PC2 included where they improved explanatory power. Prior to analysis, multivariate normality was assessed using the ‘mardia*’* function in the *mvnormalTest* R package, and variables were transformed as required. SEM analyses were conducted using the *lavaan* and *semptools* packages, with model structure and path coefficients visualised using and *semPlot*.

**Figure 3.**
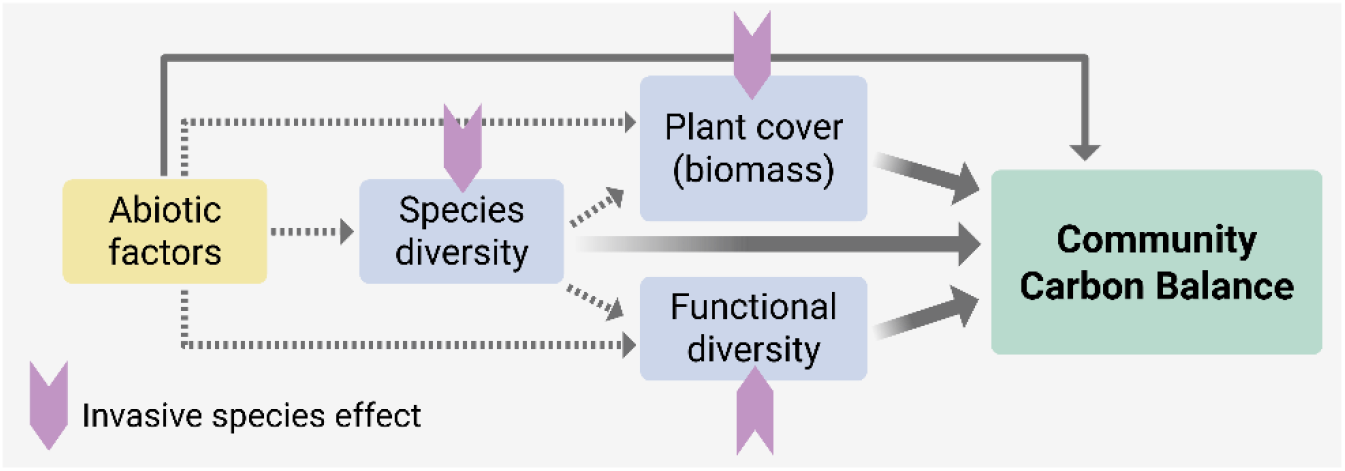
*A priori* path model (after Yan et al., 2023), illustrating hypothesised direct (solid arrows) and indirect (dashed arrows) effects on community carbon balance (NEE, GPP, ER). Invasion effects (thick grey arrows) operate through changes in diversity, biomass and community traits (pink arrows).

## Results

### Hyp_1_: Treatment effects on C fluxes

At the heather site, most treatments exhibited net zero NEE, indicating similar rates of GPP and ER (Fig. 4). However, the WR treatment significantly increased net C loss (*X*^*2*^_(1)_ = 14.31, *p* = 0.00015), with higher NEE relative to both A and AC plots (*X*^*2*^_(1)_ = 9.33, *p* = 0.010; Fig. 4). The difference in net carbon exchange between A and WR plots was 1.1 µmol CO_2_ m^-2^ s^-1^, with the WR plots acting as a net C source. Removal reduced GPP independently of warming, with AR plots showing less negative GPP than AC and WC plots (Fig. 4). In contrast, ER responded strongly to warming, increasing by 0.93 µmol CO_2_ m^-2^ s^-1^ in WC plots and 1.27 µmol CO_2_ m^-2^ s^-1^ in WR plots relative to A (Table S1). At the hawkweed site, no treatment effects on C fluxes were detected after two years (Fig 4). All treatments maintained a net zero NEE despite that GPP and ER differed significantly from zero (*X*^*2*^_(1)_ = 98.1, *p* < 0.001 and *X*^*2*^_(1)_ = 96.9, *p* < 0.001 respectively; Table S3).

**Figure 4.**
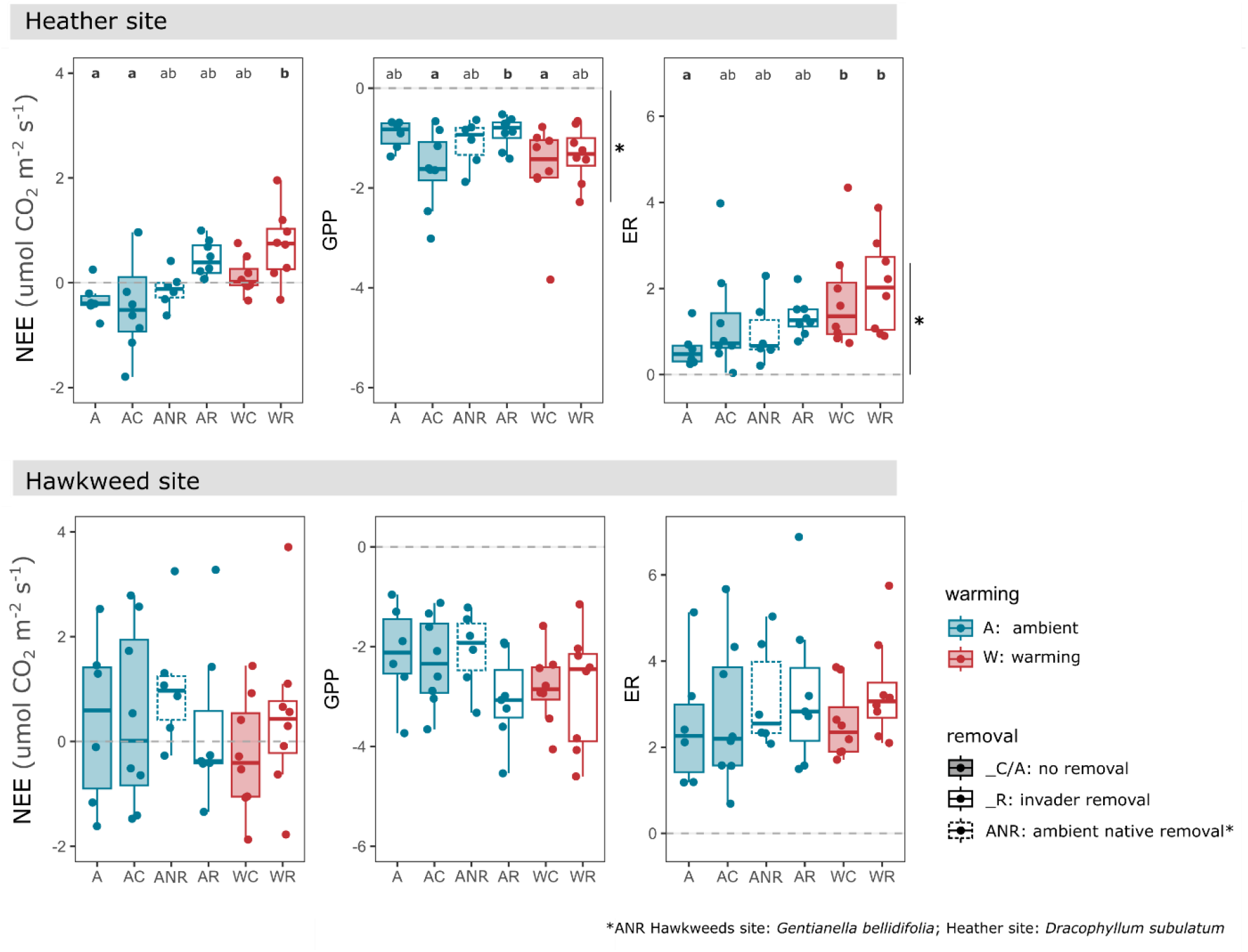
Treatment effects on carbon fluxes at the hawkweed (top) and heather (bottom) sites. Net ecosystem exchange is shown alongside its component fluxes (NEE = GPP - ER). For NEE, values below zero indicate net carbon uptake, values above zero indicate net carbon loss, and values overlapping zero indicate a neutral carbon balance. Significant differences among treatments are shown with different letters (*e*.*g*. a, b), and absence of letters denotes no significant treatment effects (e.g. Hawkweed site). All fluxes are reported in µmol CO_2_ m^-2^ s^-1^.

### Hyp_1_: Treatment effects on abiotic and biotic context

#### Heather site

Mean air temperatures under warming were ∼1 °C higher than in ambient plots, while mean maximum air temperatures increased by 2-5 °C (*X*^*2*^_(1)_ = 43.56, *p* = 0.0023, Fig. S1a). Dominant plant removal had no effect on air temperature (*X*^*2*^_(1)_ = 50.00, *p* < 0.0001). Soil moisture was significantly lower in A than in WR plots (*X*^*2*^_(1)_ = 17.25, *p* = 0.016, Fig. S1b). Total plant cover was highest in A plots (*X*^*2*^_(1)_ = 64.3, *p* = 0.010), where *C. vulgaris* is naturally absent (Fig. S1c). Species richness did not differ significantly among treatments, although plots were *C. vulgaris* was absent or removed (A, AR, WR) tended to have slightly higher species richness (Fig. S1d). While removal had no overall effect on richness, invader cover was negatively correlated to species richness when invaded plots were considered alone (*X*^*2*^_(1)_ = 22.54, *p* = 2.05e^-6^). In ambient plots, *C. vulgaris* cover increased by approximately 10% between 2015 and 2022 (*t* = - 3.37, *p* < 0.01), with no significant effect of warming on invader cover.

The first two axes of a PCA explained 89% of the variation in plant traits, biomass and diversity (Fig. S2). PC1 was strongly associated with invader dominance, with major contributions from *C. vulgaris* cover, CWM leaf thickness, and NDVI (*cor*: 0.94, *p* = 1.03e^-21^; *cor*: 0.74, *p* = 7.15e^-09^; *cor*: -0.92, *p* = 2.61e^-19^ respectively). PC1 clearly separated treatments invaded from uninvaded or removal plots (i.e., A, AR and WR; Fig S2). In contrast, PC2 was primarily composed of species evenness (*cor*: 0.91, *p* = 1.47e^-18^) and greenness (*cor*: 0.43, *p* = 3.41e^-03^) and did not correspond to treatment effects. Invader cover was positively correlated with total biomass (*cor* = 0.8; *p* < 0.001, FS4) and NDVI (*cor* = 0.6; *p* < 0.001), but negatively correlated with species richness (*cor* = -0.64; *p* < 0.001), Shannon’s diversity (*cor* = -0.45; *p* < 0.01), evenness (*cor* = -0.17; *p* < 0.05), and cover of the subdominant shrub *D. subulatum* (*cor* = -0.56; *p* < 0.01). Invader cover was also negatively correlated with key CWM traits, including leaf thickness (*cor* = -0.63; *p* < 0.001), δ^13^C (more negative, *cor* = 0.44; *p* < 0.001) and SLA (*cor* = - 0.61; *p* < 0.001), indicating strong shifts in community structure and trait composition with invasion.

#### Hawkweed site

Warming increased mean air temperature by ∼1°C (Fig S1a) and elevated maximum air temperatures by 6 -10 °C relative to ambient plots (*X*^*2*^(1) = 43.70, *p* = 0.003). Soil moisture did not differ significantly among treatments (*X*^*2*^(1) = 3.46, *p* < 0.001), although WR plots tended to have the lowest values (Fig S1b). Total plant cover and species richness were higher in WC than in A and ANR plots (Fig S1g: *X*^*2*^(1) = 4.54, *p* = 0.033; Fig S1h: *X*^*2*^(1) = 5.64, *p* = 0.017, respectively). However, invasive hawkweed cover declined by 7.8% under warming (*X*^*2*^(1) = 4.88, *p* = 0.027).

The first PCA axes explained 74.4% of the variation in plant traits, biomass and diversity (Fig S3). Both axes were driven primarily by biomass and diversity-related variables, with minimal contribution from abiotic variables and trait diversity (FD_Q_). PC1 was positively associated with NDVI (*cor*: 0.81; *p* = 2.05e^-11^) and plant cover (*cor* = 0.76; *p* = 6.20e^-12^) and a negatively associated with CWM δ^13^C (more negative, *cor*: -0.64; *p* = 2.59e^-6^). PC2 was mainly driven by species richness (*cor*: 0.72; *p* = 3.31e^-8^) and CWM leaf respiration (Rd; *cor*: -0.74, *p* = 1.37e^-8^). Treatments had little effect on on multivariate community composition, as indicated by weak clustering in PCA space (Fig S3). Nevertheless, hawkweed cover was negatively correlated with species richness (*cor* = -0.47; *p* < 0.001) and total plant cover (Fig S4b; *cor* = -0.43; *p* < 0.05), and CWM leaf respiration (*cor* = -0.39; *p* < 0.01), and positively correlated with CWM δ^13^C (more negative; *cor* = 0.59; *p* < 0.01), suggesting the invasive forb reduced plot-level leaf respiration and drought resistance.

### Hyp_2_: Drivers of C fluxes – linear models

At the heather site, linear mixed-effects models showed that warming increased both GPP (*X*^*2*^_(1)_ = 9.22, *p* = 0.0023) and ER (*X*^*2*^_(1)_ = 31.14, *p* = 0.043), resulting in greater net C loss (NEE > 0; Table S2). Total plant cover was a strong positive predictor of all fluxes (NEE: *X*^*2*^_(1)_ = 5.93, *p* = 0.014; GPP: *X*^*2*^_(1)_ = 8.83, *p* = 0.0029; ER: *X*^*2*^_(1)_ = 5.72, *p* = 0.016). Likewise, PC1 representing invader biomass and associated traits, had a significant positive effect on GPP (*X*^*2*^_(1)_ = 9.57, *p* = 0.0019; Table S2), indicating that higher *C. vulgaris* dominance increased C uptake (Fig. S2). This was further supported by a positive relationship between GPP and invader cover (*p* = 0.081; R^2^ = 0.39; Fig. S6).

At the hawkweed site, both PC1 (biomass/NDVI) and PC2 (species richness and leaf respiration) influenced C fluxes (Table S4). PC2, was a significant predictor of NEE (*t* = -3.03, *p* = 0.005), while PC1 had a marginal effect (*t* = -1.86, *p* = 0.071). Similarly, both PC1 and PC2 had a significant effect on GPP (PC1: *t* = 4.07, *p* < 0.0001; PC2: *t* = 2.27, *p* = 0.030). GPP significantly increased with NDVI (*R*^*2*^: 0.54, *p* < 0.001), species richness (*R*^*2*^: 0.37; *p* = 0.016), and total plant cover, especially under removal (*R*^*2*^: 0.55, *p* = 0.01; Fig S7). Hawkweed removal had a significant positive effect on ER (*t* = -2.54, *p* = 0.016), while plots with higher species evenness (*t* = -2.48, *p* = 0.018) and CWM RWC (*t* = -2.82, *p* = 0.008) respired less (Table S4).

### HYP_2_: Drivers of C fluxes – SEM

#### Heather site

The SEM explained 22% of the variation in NEE at the heather site. Air temperature had a direct positive effect on NEE (0.26, *p* = 0.034), increasing net C loss (Fig. 5). Species diversity also strongly influenced NEE. Species richness had a direct positive effect on NEE (0.29, *p* = 0.033) but also an indirect negative effect mediated through reduced NDVI (-0.33, *p* = 0.004), which in turn decreased NEE (-0.31, *p* = 0.05). This indicates that uninvaded, species rich plots were less green (lower NDVI) and had lower C uptake than heather-invaded plots. Higher species richness was also associated with greater CWM leaf thickness (0.57, *p* < 0.0001), which was negatively related to soil moisture, such that plants in drier plots had thicker leaves (L_th_, -0.23, *p* = 0.019; Fig. 5).

**Figure 5.**
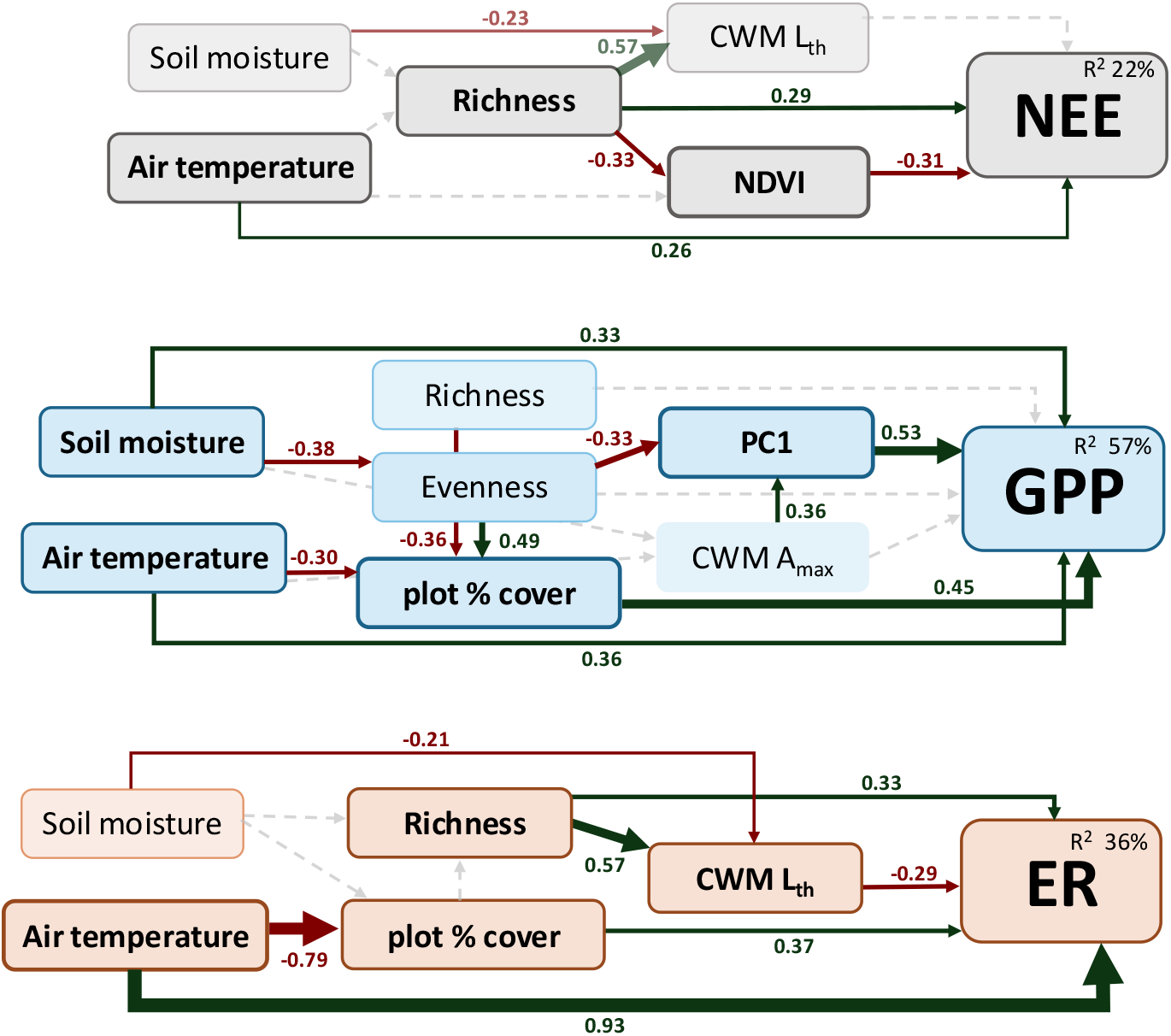
Structural equation models (SEM) with path analysis for NEE, GPP and ER at the heather site, based on the *a priori* model. Arrows indicate relationships included in the final model: solid arrows denote significant paths, with green and red indicating positive and negative effects, respectively, while dashed arrows indicate non-significant relationships. Arrow width is proportional to effect size, and standardised path coefficients are shown adjacent to each arrow. The proportion of variance explained (R^2^) for each flux is shown within the response variable box. Variables with outlined boxes have significant direct effects on the respective flux. PC1 primarily represents *C. vulgaris* biomass and associated traits.

The SEM explained 57% of the variation in GPP. PC1, representing invader biomass and traits, had the strongest positive effect on GPP (0.53, *p* < 0.0001). Abiotic factors also contributed, with with both air temperature (0.36, *p* = 0.002) and soil moisture (0.33, *p* = 0.004) increasing GPP. Plant cover was also positively associated with GPP (0.46, *p* = 0.001). Species evenness had an indirect negative effect on GPP through PC1 (-0.33, *p* = 0.008), indicating that plots dominated by *C. vulgaris* had lower evenness.

The SEM explained 36% of the variation in ER. Air temperature exerted the strongest direct positive effect (0.93, *p* < 0.001), indicating higher respiration in warmer plots. Soil moisture had an indirect negative effect on ER (-0.21, *p* = 0.023) through its influence on leaf thickness, with thicker leaves reducing ER (-0.29, *p* = 0.034). Plant cover and species richness also increased ER (0.37, *p* = 0.035, 0.31, *p* = 0.032, respectively). However, the positive direct effect of species richness (0.33, *p* = 0.020) was offset by its indirect negative effect on leaf thickness, which reduced ER, resulting in lower C efflux in species-rich plots (0.57, *p* < 0.0001).

#### Hawkweed site

The SEM explained 26% of the variation in NEE at the hawkweed site (Fig 6). In contrast to the heather site, abiotic variables had no direct effects on NEE. Instead, leaf traits were the primary drivers: increases in CWM leaf thickness (-0.28, *p* = 0.003) and leaf respiration (-0.29, *p* = 0.015) directly enhanced C uptake (more negative NEE). Plant biomass had a direct positive effect on NEE (-0.24, *p* = 0.025), indicating greater C loss with increasing biomass. Species richness indirectly increased C uptake through a positive effect on CWM leaf respiration (0.40, *p* = 0.001), highlighting the importance of diversity-mediated trait pathways.

**Figure 6.**
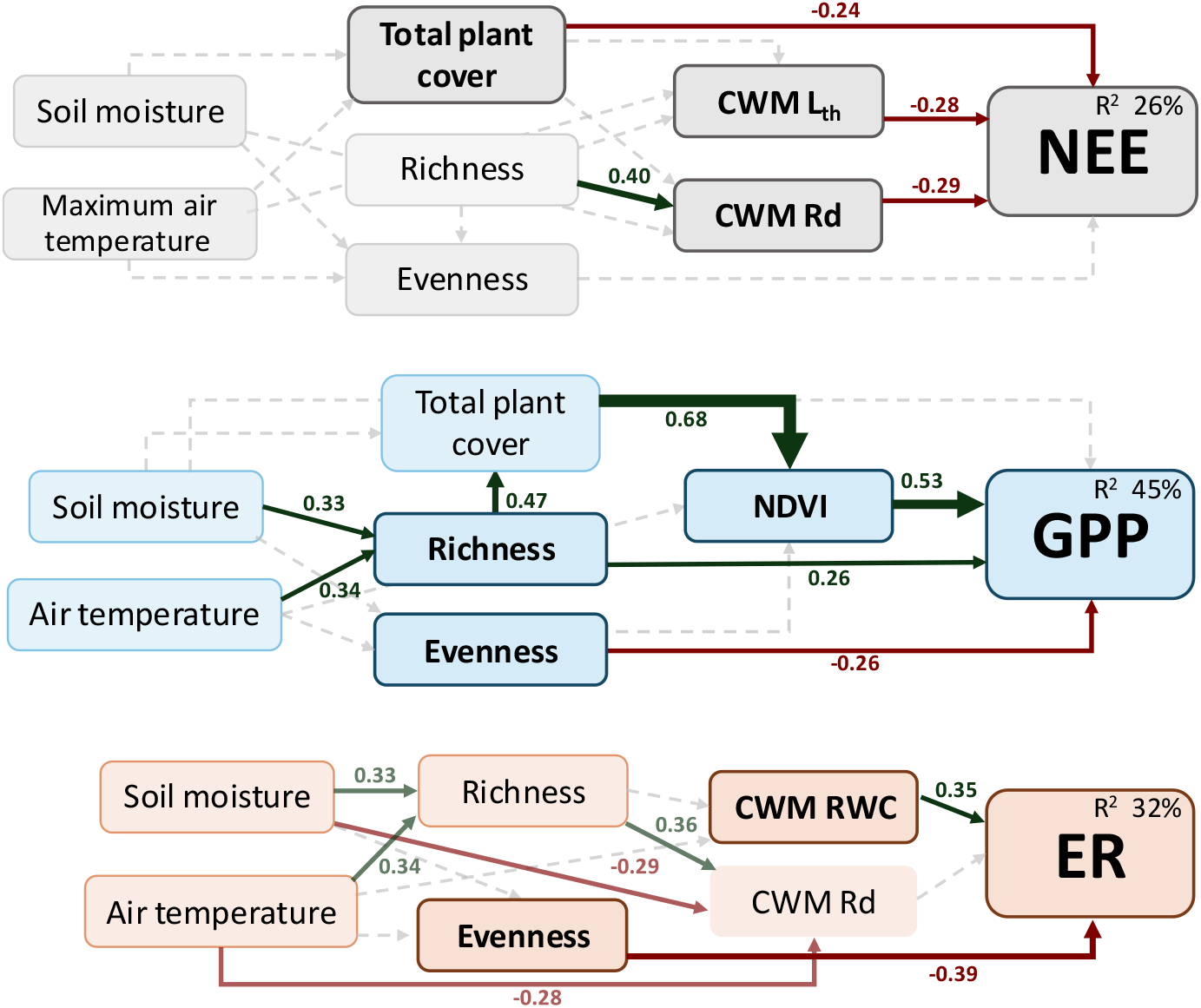
Structural equation models (SEM) with path analysis for NEE, GPP and ER at the hawkweed site, based on the *a priori* model. Arrows indicate relationships included in the final model: solid arrows denote significant paths, with green and red indicating positive and negative effects, respectively, while dashed arrows indicate non-significant relationships. Arrow width is proportional to effect size, and standardised path coefficients are shown adjacent to each arrow. The proportion of variance explained (R^2^) for each flux is shown within the response variable box. Variables with outlined boxes have significant direct effects on the respective flux.

The SEM explained 45% of the variation in GPP. NDVI and species richness both had direct positive effects on productivity (0.53, *p* < 0.0001; 0.26, *p* = 0.047), while evenness had a negative effect (-0.26, *p* = 0.013). Total plant cover indirectly increased GPP through its positive effect on NDVI (0.68, *p* < 0.0001). Abiotic variables (air temperature and soil moisture) only affect GPP indirectly through their positive effects on species richness, further supporting a diversity-driven mechanism of productivity.

Finally, the SEM for ER explained 32% of total variance. Species evenness had a direct negative effect on ER (-0.39, *p* = 0.003), while CWM RWC showed a marginal positive effect (0.35, *p* = 0.055), indicating higher C efflux in plots with greater leaf turgidity and lower evenness. No significant relationship was detected between CWM leaf dark respiration and ER (*p* = 0.118). Instead, CWM R_D_ was best explained by abiotic conditions and species diversity: drier soils and cooler temperatures increased leaf respiration, as did species richness (0.36, *p* = 0.016). Consistent with the absence of direct abiotic controls on ER, soil moisture influenced fluxes indirectly via its negative relationship with CWM leaf respiration, such that plots with higher water availability had lower leaf-level respiration at the hawkweed site (Fig 6).

## Discussion

### Treatment effects on carbon balance

Overall, we found no evidence of positive feedback between warming and plant invasion. At the heather site, warming offset invasion-driven productivity gains by increasing ecosystem respiration, resulting in net carbon loss under combined warming and removal (WR). Contrary to our first hypothesis, warming did not enhance net carbon uptake: it had no effect at the hawkweed site and drove carbon loss at the heather site. This outcome arose from reduced gross primary productivity (GPP) following removal and increased ER under warming. While invader biomass and traits supported higher productivity, higher temperatures and plant cover were associated with increased respiration, shifting the balance towards carbon loss. These results indicate that removing the invasive shrub can reduce ecosystem carbon uptake, particularly under warming. Consistent with this, previous work at the heather site showed warming increased GPP only in non-invaded plots (Vought et al., 2026), aligning with the net carbon uptake we observed in uninvaded communities. Together with long-term evidence that *Calluna vulgaris* expansion occurs primarily under ambient conditions (Leon-Garcia et al., 2025), these findings suggest that warming-invasion feedbacks are constrained, likely by factors such as low water availability (Jiao et al., 2021).

Under experimental warming, ER showed the strongest response, leading to net carbon loss in the WC and WR treatments. This contrasts with meta-analyses indicating that GPP is typically more sensitive to warming, resulting in net carbon gains in colder grasslands (Chen et al., 2017; Wang et al., 2019). In our system, the absence of a GPP response likely reflects constraints on productivity under warming. Reduced plant cover in warmed plots (Leon-Garcia et al., 2025) and shifts in resource availability, particularly water limitation, may have suppressed photosynthetic gains (Harte, Scott & Charlotte, 2015; Yang et al., 2018). Previous work at these sites shows that *C. vulgaris* reduces community-level drought tolerance by lowering leaf thickness and water content, reinforcing water limitation as a key constraint on ecosystem functioning (Leon-Garcia & Deslippe, 2025). Warming-induced changes in community composition, including increases in drought- and low-nutrient-tolerant plant species (Leon-Garcia et al., 2025; Patton et al., 2007), may further offset any potential productivity increases (Fu & Shen, 2017; Wang et al., 2020). In addition, warming can exacerbate heat and moisture stress, reducing microbial activity and nutrient availability (Maestre et al., 2015; Sardans et al., 2008). These mechanisms are consistent with experimental studies in alpine systems showing limited or absent GPP responses to warming, but strong sensitivity to water availability. For example, soil moisture strongly mediates GPP responses in alpine meadows in China (Fu et al., 2018; Zhang et al., 2024). Together, our results reinforce the view that water availability constrains warming responses in cold, dry ecosystems, limiting gains in productivity despite increased temperature (Lyu et al., 2026; Yang et al., 2018).

Removal of hawkweed increased ecosystem respiration but had no effect on net carbon balance. Contrary to expectations removal did not enhance carbon uptake, but instead elevated respiration, regardless of whether native or invasive species were removed, and this effect persisted even after temperature correction. While removal had no direct effect on GPP, total plant cover was positively associated with GPP, with this relationship strengthening under removal (Fig. S7). The increase in ER may therefore reflect tighter coupling between photosynthesis and respiration, as higher productivity is often accompanied by increased respiratory demand (Flexas et al., 2007). Alternatively, the increase in respiration following removal may result from increased decomposition of belowground root and stolon biomass, and greater bare ground exposure following removal (Wan et al., 2005; Zhu et al., 2024), as well as short-term disturbance effects from treatment application given the relatively short duration of the experiment at this site.

Treatment effects were pronounced at the heather site but weak at the hawkweeds site, likely reflecting the shorter treatment duration in slower-growing, higher-elevation communities. Although warming rates were similar, longer treatment exposure at the heather site likely allowed greater shifts in plant biomass and community composition. In contrast, the hawkweed appears less functionally distinct from native species and contributes little to plot biomass, potentially limiting its impact on carbon balance (Richardson & Pyšek, 2012). These results highlight that invasion effects on ecosystem carbon dynamics are not generalisable across invader types but depend on growth form and trait differences. Continued monitoring will be essential to determine whether these systems maintain functional stability or whether longer-term declines in diversity ultimately offset short-term productivity gains.

### Invasion effects on biodiversity and community carbon flux

Consistent with our second hypothesis, the relative importance of biodiversity and biomass differed between sites. At the hawkweed site, carbon fluxes were primarily structured by species diversity and trait composition, whereas at the heather site they were driven by biomass and invader dominance. Across both systems, water availability emerged as a key constraint, with soil moisture influencing both community composition and carbon fluxes. Together, these results indicate that the mechanisms linking biodiversity to ecosystem carbon dynamics depend strongly on invader identity, community structure, and abiotic context.

At the heather site, we observed a clear dominance-driven productivity effect: increasing *C. vulgaris* cover enhanced aboveground productivity (GPP, NDVI, biomass) while reducing native species diversity. This reflects a shift towards invader-driven biomass accumulation at the expense of community diversity, including reduced cover of the dominant native shrub *Dracophyllum subulatum*. Long-term removal experiments show that diversity gains following invader removal are primarily due to increases in small, low-biomass species, which do not compensate for the loss of productivity (Leon-Garcia et al., 2025; Livingstone et al., 2020). These results support the dominance-driven productivity hypothesis (Flombaum, 2017), whereby invasion increases productivity through functional dominance while reducing diversity. However, this apparent gain in function is likely context dependent. Dense *C. vulgaris* stands reduce the cover of moisture-dependent species (Moyle & Deslippe, 2024) and and the associated shifts in community traits we observe, including reduced leaf thickness, RWC and δ^13^C, suggest lower community drought tolerance with invasion. More broadly gains in a single function may come at the expense of others, for example, through increasing fire risk (Schirmel et al., 2016; Tripathi et al., 2023). Together, these findings indicate that productivity gains associated with invasion may be transient, with longer-term negative consequences for biodiversity and ecosystem functioning.

In contrast, carbon dynamics at the hawkweed site were driven primarily by diversity rather than invader dominance. The invasive forb had minimal direct effects on carbon fluxes, instead influencing them indirectly through reductions in species richness and native cover. Soil moisture was positively correlated to species richness, again highlighting the sensitivity of community composition to water availability. Hawkweed abundance was correlated with lower species richness and occurred preferentially in more open vegetation, consistent with its affinity for less densely vegetated grasslands (Steer & Norton, 2013). Unlike the heather system, where invasion increased productivity through biomass dominance, carbon uptake at the hawkweed site was positively associated with species richness, total plant cover and leaf respiration. Consequently, hawkweed invasion may reduce carbon uptake indirectly via its negative effects on diversity. However, given its low contribution to biomass and limited spread, invasion appears unlikely to substantially alter ecosystem carbon balance under current conditions.

While our experimental design captures key aboveground drivers of carbon flux, it likely underestimates the contribution of belowground processes, particularly to ecosystem respiration, which integrates both autotrophic and heterotrophic components. The stronger explanatory power of GPP relative to ER in our models reflects this complexity and highlights the need to better resolve belowground controls on carbon balance. Across both sites, biotic variables, especially plant cover, NDVI, and species diversity, emerged as primary drivers of carbon uptake, but these relationships were consistently constrained by water availability. Collectively, our results show that the balance between biodiversity and biomass in regulating carbon fluxes is context dependent, shaped by invader identity and modulated by water-stress. Future work integrating belowground traits, soil microbial dynamics, and longer-term responses to warming and disturbance will be critical for predicting whether invasion-driven gains in productivity persist or give way to reduced ecosystem resilience under increasing climatic stress.

