## Supplementary material for "Community and leaf-level controls of carbon fluxes in an invaded NZ alpine tussock grassland"

**Additional methods.** *NEE chamber design and calculations*

We used a custom-made 0.7 $m^3$ (1m×1m×0.7m) clear plexiglass chamber with full spectrum artificial light from five dimmable lamps (Figure S0), to calculate *in situ* plot level $CO_2$ fluxes (net ecosystem exchange, NEE; gross primary productivity, GPP; and ecosystem respiration, ER). We partitioned the NEE into carbon influx (GPP) and efflux (ER; Chapin et al., 2006), by recording $CO_2$ exchange in darkness to estimate ER, and calculating GPP as:

$$NEE = GPP - ER$$

the obtained ER under darkness encompasses both soil heterotrophic and above- and below-ground plant respiration (*i.e.* autotrophic respiration, Chapin et al., 2006). We recorded net carbon exchange with a LI-COR 7500A (LI-COR, Inc., Lincoln, NE, USA) inside the polycarbonate chamber in a closed system at peak productivity (Jan-Feb). We conserved the notation LI-COR uses for recorded $CO_2$ absorptance, were GPP is shown as a negative value, signifying the decrease in $CO_2$ concentration when net C uptake occurs (NEE < 0), and ER is a positive value reflecting net C loss as the $CO_2$ concentration increases inside the enclosed chamber with respiration (NEE > 0). We prevented leaks and ensured the chamber was airtight by installing two tarpaulin skirts weighed to the ground with two sets of chains for a tight close even on uneven ground. The air inside the enclosed chamber was mixed with four fans. We recorded air temperature and light inside at two points under the lamps and at canopy level (~60 cm) during the measurements. We standardize carbon exchange at one light point (*e.g.* $NEE_{800}$) to allow among plots and sites comparisons. In order to calculate this, we measured C fluxes in darkness and at three light levels, achieved with dimmable lights set at 0%, 50%, 75% and 100% capacity in that order. We powered the lights with a generator. We first recorded ecosystem respiration (ER) measuring $CO_2$ exchange in darkness, setting off the lamps (0%) and with a dark cloth over the transparent chamber. The three light points followed. Photosynthetic active radiation (PAR) under the lamps for the three light points ranged from 200 to 3000 μmol $m^{-2}$ $s^{-1}$ of PAR under 100%. We ventilated the chamber in between measurements to allow carbon and air temperature to stabilized and return to ambient values before starting the next recording. We collected data for one to two minutes after the chamber was closed and carbon reached stability. All NEE measures were recorded between 8 am and 4 pm on sunny days and it took around two weeks to complete each site. We recorded soil moisture with a Delta T Devices HH2 Moisture Meter each measuring day to ensure water availability didn't change, especially after rainy days. When soil conditions varied too much, we waited a day for soil moisture to return to the set standard value at previous days. Net Ecosystem Carbon Exchange (NEE) for each curve was calculated according to Prager et al., 2021:

$$\rho = \frac{P}{8.314 \times T}$$

$$NEE = (\rho \times V \times (\frac{dC}{dt})/A)$$

were $\rho$ was the air density, calculated with the mean pressure $P$ (Pa), the ideal gas constant, and the mean air temperature $T$ (K) extracted from the LI-COR at the time of measure. For the NEE equation, $V$ was the chamber volume, and $dC/dt$ the slope of the $CO_2$ absorptance against time curve, and $A$ the surface ground area inside the chamber (1$m^2$) (Prager et al., 2021). We calculated this value for each light point and under darkness, were GPP is zero so NEE equals ER. To perform across treatment comparisons, we used the light response curve (LRC) of NEE, to standardize C fluxes to a single light value per plot (800 µmol $m^{-2}$ $s^{-1}$) using the equation from this relation to calculate $NEE_{800}$ and $GPP_{800}$ (De Lobo et al., 2013; Sundqvist et al., 2020) as follows:

$$NEE_{800} = intercept_{LRC} + slope_{LRC} \times 800\ PAR$$

$$ER = intercept_{LRC}$$

$$GPP_{800} = NEE_{800} - ER$$

All presented values of NEE and GPP and ER in this paper correspond to this standardized value at 800 PAR, unless stated otherwise.

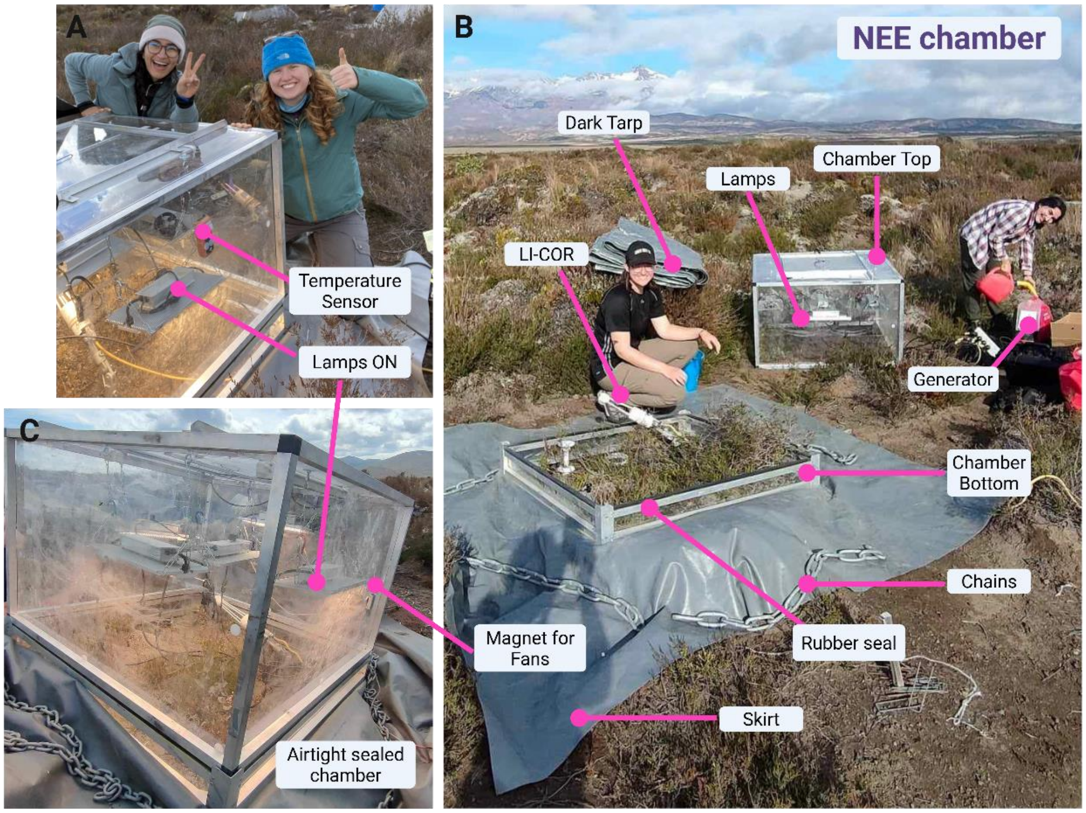


**Figure S0.** NEE chamber design and labelled parts. A) and C) show the complete assembled NEE chamber. B) Shows the chamber top and bottom parts of the NEE chamber. A clear box was chosen instead of an opaque one to facilitate troubleshooting during measurements (i.e. branches on laser path for the LI-COR, fans working correctly… etc). All measurements (including darkness and the three light points) were conducted under a dark tarp to ensure that recordings were made without any ambient light. Photo C) shows the lights ON without the tarp just for illustrative purposes. The generator was used to power the five lamps and the computer. We powered the LI-COR 7500A with a 12V battery.

**Figures.**

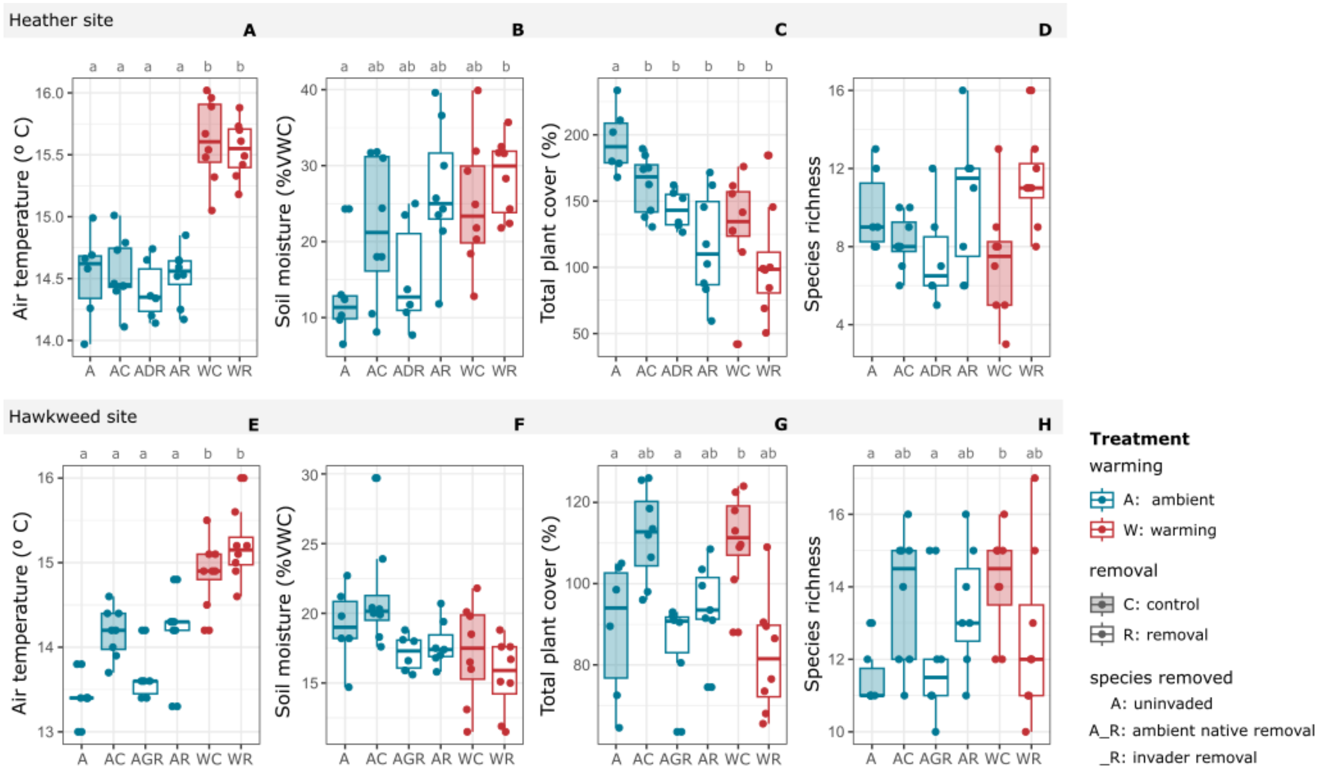


**Figure S1.** Treatment effects on environmental conditions, plant cover and diversity at both sites. A-D corresponds to the Heather site and E-H to the Hawkweed site. Blue plots correspond to ambient plots while red depicts the warming treatments. Statistical difference among treatments is represented with different letters, no letters mean no significant treatment effects were found.

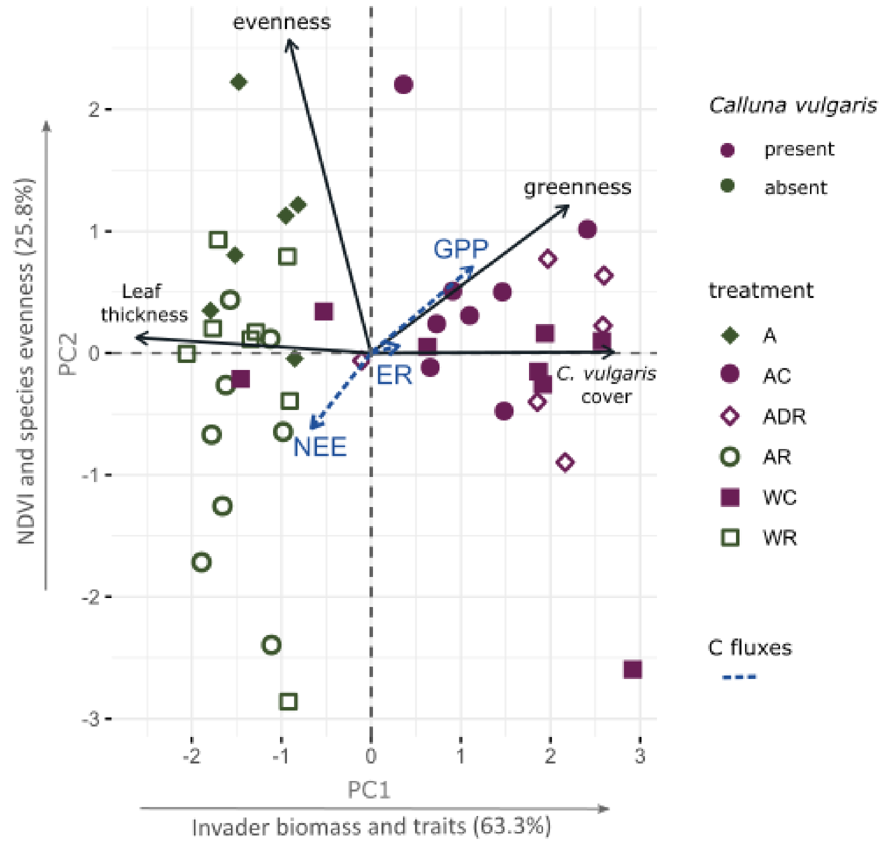


**Figure S2.** A PCA of the CWM of plant traits, biomass estimates and species diversity indexes for the plots at the invasive heather site, explaining a cumulative variance of 89.1%. Black vectors length and direction indicates the strength of their correlation with PC1 and PC2. Dashed blue vectors correspond to C fluxes, included in the biplot as supplementary quantitative variables. Closeness to the black predictors correspond to high correlation with this variables. Hollow dots correspond to removal plots (i.e. ADR, AR, WR) and filled dots to non-removal plots. Purple treatments correspond to *C. vulgaris* invaded plots and the green dots to uninvaded ones. GPP was multiplied by -1 to show both gross productivity and ecosystem respiration as positive values for an easier graph interpretation.

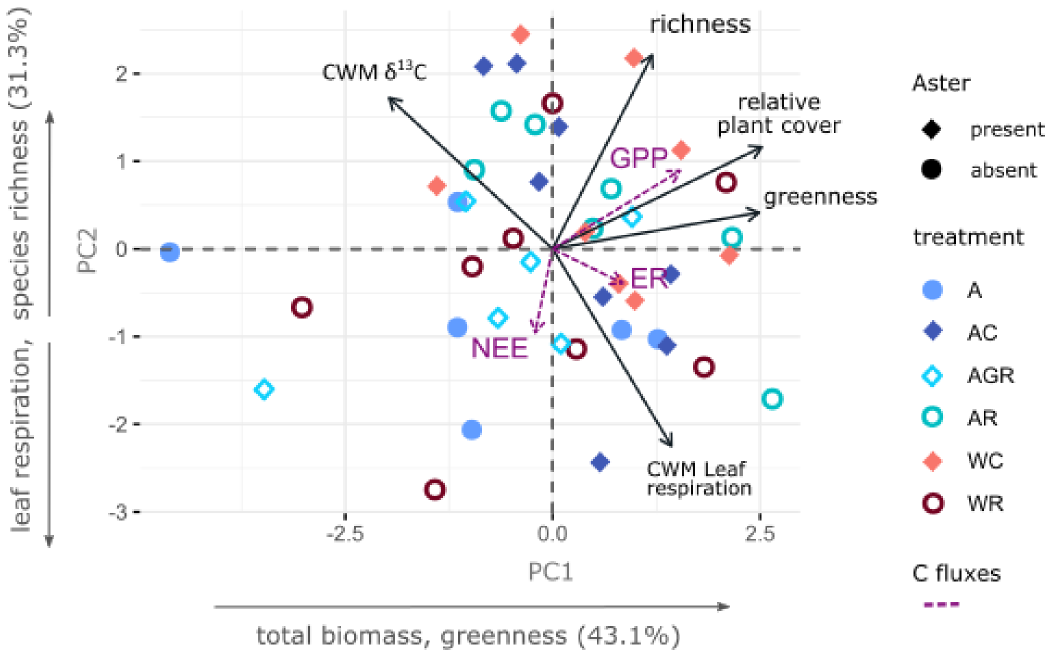


**Figure S3.** A PCA for the CWM of plant traits, biomass estimates and species diversity indexes for the plots at the invasive hawkweed site, cumulative variance explained is 74.4%. Dashed vectors correspond to C fluxes, included to the biplot as supplementary quantitative variables. Hollow dots correspond to removal plots (i.e. AGR, AR, WR) and filled dots to non-removal plots.

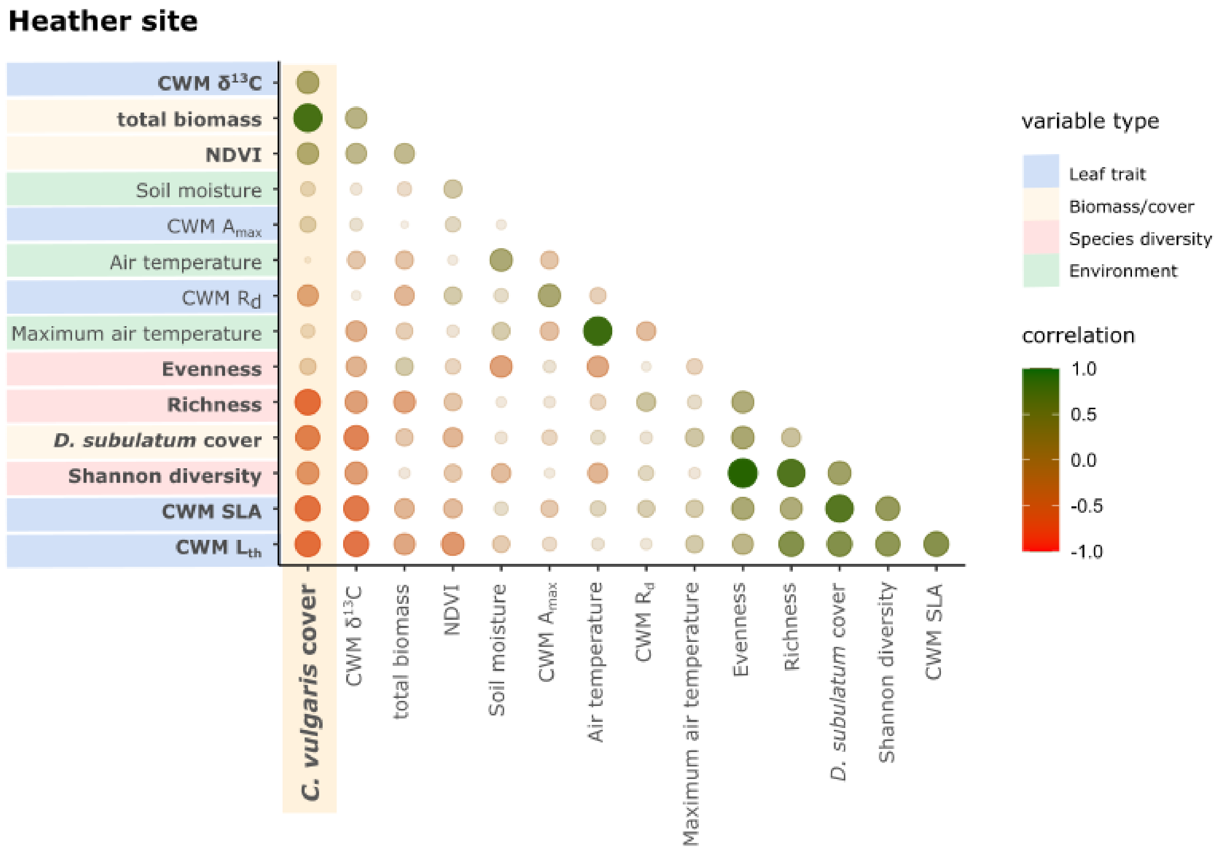
Heather site
CWM δ13C
total biomass
NDVI
Soil moisture
CWM Amax
Air temperature
CWM Rd
Maximum air temperature
Evenness
Richness
D. subulatum cover
Shannon diversity
CWM SLA
CWM Lth
C. vulgaris cover
variable type
Leaf trait
Biomass/cover
Species diversity
Environment
correlation
1.0
0.5
0.0
-0.5
-1.0

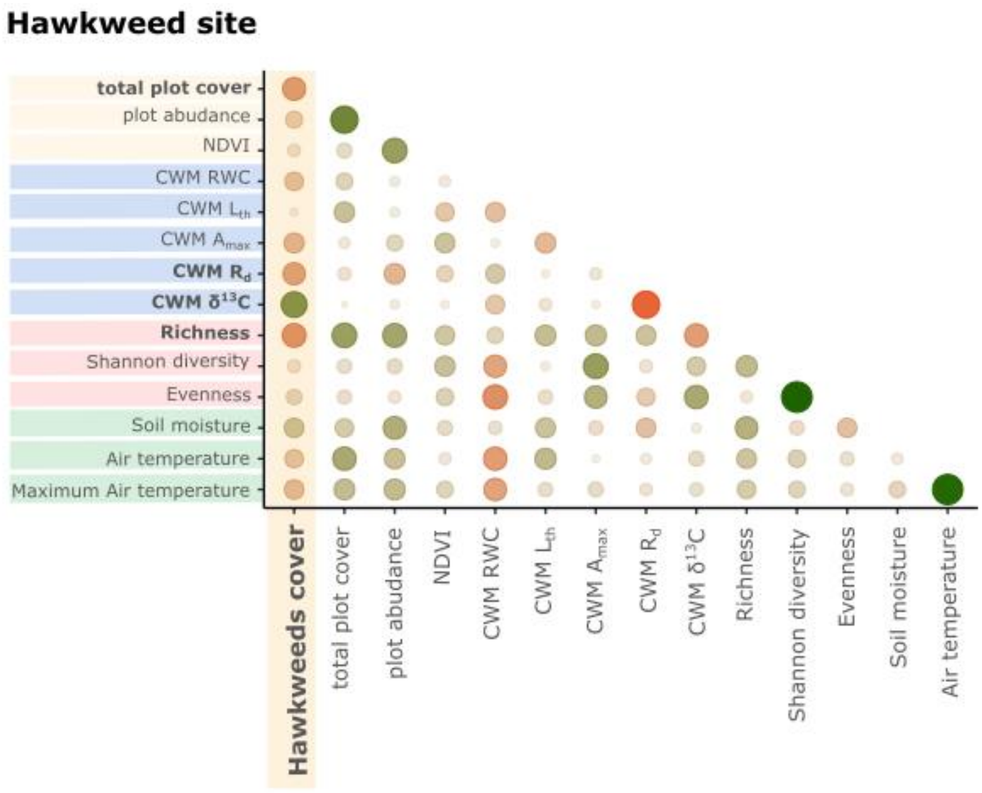
Hawkweed site
total plot cover
plot abudance
NDVI
CWM RWC
CWM Lth
CWM Amax
CWM Rd
CWM δ13C
Richness
Shannon diversity
Evenness
Soil moisture
Air temperature
Maximum Air temperature
Hawkweeds cover

**Figure S4. Correlation matrix for invader percent cover and all predictors at both sites.** Matrix for heather site (**top**) and hawkweed site (**bottom**). Variables are classified according to their type and this is depicted by the different colors: community weighted mean (CWM) of leaf trait (blue), biomass or cover (yellow), species diversity index (red) or environment (green). CWM of $R_d$ and $\delta^{13}C$ were transformed to positive values for an easier interpretation, so positive correlation corresponds to an increase in the number value of these traits (*i.e.* more negative). The strength of the correlations is represented by the size of the circle and positive correlations are shown in green and negative in red. Names shown in bold depict statistically significant correlations to invader cover. The significance among other predictors is not shown but correlation strength corresponds to dot size and color intensity.

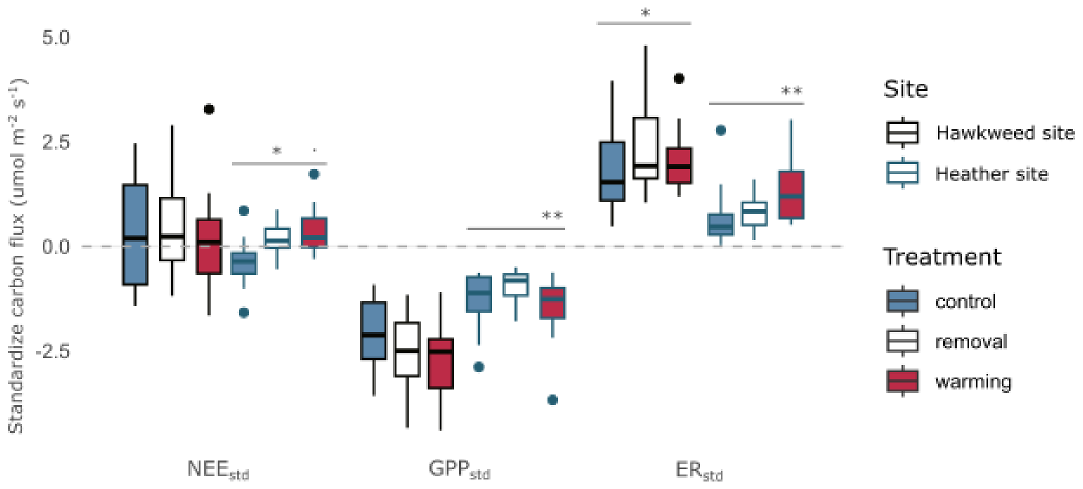


**Figure S5. Treatment effects on standardized carbon fluxes for both sites.** Fill color depicts treatments grouped in removal (AR, WR), warming (WC, WR) and control (A, AC, ANR). Border color corresponds to the site. Net carbon balance is depicted by the dashed line in the zero value. All statistically significant differences shown were calculated comparing to the control (blue). Each flux was standardized by the standard deviation of the C fluxes at both sites. The number of asterisks denote the significance: '**' when $p < 0.01$, '*' when $p < 0.05$ and '.' when $p < 0.1$.

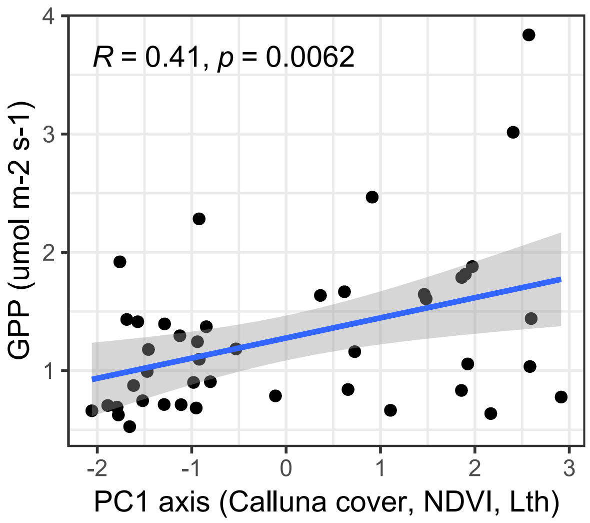


**Figure S6. Linear regression model for GPP and the PC1 axis at the heather site.** PC1 axis has the biggest contribution from *C. vulgaris* cover, NDVI and leaf thickness. Model statistics are included for the fitted line. Gray area is the confidence interval of 95% of the model.

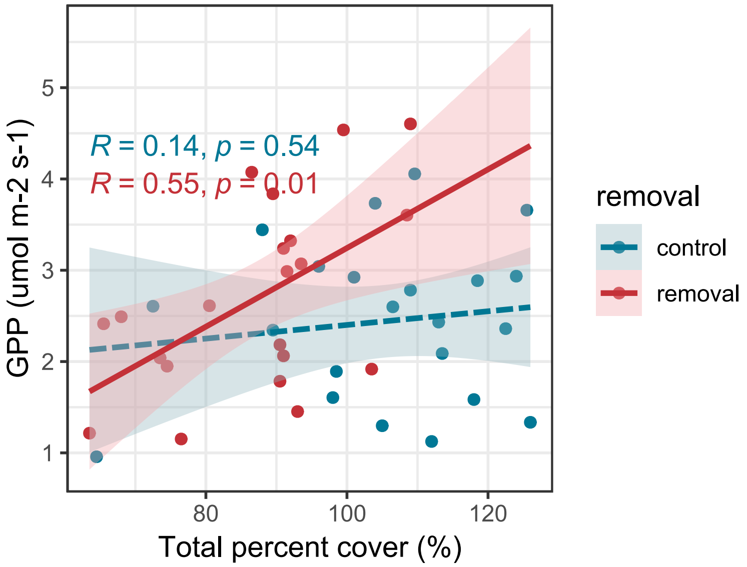


**Figure S7. Linear model for GPP and total percent cover at the hawkweed site.** Models were fit by invader removal. Colored area is the confidence interval of 95% of each model. Model statistics are included for each fitted line. Dashed line for model fit corresponds to the lack of significance. After hawkweed removal, GPP responded more to total plot cover.

**Tables.**

**Table S1.** Linear mixed effect models summary table for the treatment effects (warming: W, removal: R) on C fluxes separately (NEE, ER, GPP) for the heather site. The A treatment corresponds to uninvaded ambient plots, while AC are ambient and invaded plots. The number of asterisks denote the significance: ** when $p < 0.01$ and * when $p < 0.05$.

| | Heather site - **Treatment effects** | | | | |
|---|---|---|---|---|---|
| C flux | treatment contrast | *ratio* | *t* ratio | *p* value | significance |
| NEE | **A - WR** | -1.11 | -2.90 | **0.007** | ** |
| | **AC - WR** | -1.11 | -3.52 | **0.001** | ** |
| GPP | **AC - AR** | 1.95 | 2.99 | **0.005** | ** |
| | **AR - WC** | 0.56 | -2.83 | **0.008** | ** |
| ER | **A - WC** | 0.63 | -2.28 | **0.028** | * |
| | **A - WR** | 0.55 | -2.62 | **0.013** | * |

**Table S2.** Linear mixed effect models summary table for the treatments and predictors of C fluxes separately (NEE, ER, GPP) for the heather site. The number of asterisks denote the significance: '***' when $p < 0.001$, '**' when $p < 0.01$, '*' when $p < 0.05$ and '.' when $p < 0.1$. PC1 and PC2 were the two principal components extracted from the PCA analysis. PC1 summarizes the invaders biomass and traits, while PC2 has the most contribution from species diversity.

| | | Heather site | | | |
|---|---|---|---|---|---|
| **C flux** | **Predictor** | **Estimate** | ***t* value** | **Pr(>\|*t*\|)** | **significance** |
| **NEE** | warming | -0.35 | -1.76 | 0.087 | . |
| | **removal** | -0.24 | -2.30 | **0.027** | * |
| | warming : removal | 0.01 | 0.11 | 0.917 | |
| | air temperature | 0.41 | 1.04 | 0.307 | |
| | **max air temp** | -0.12 | -2.19 | **0.035** | * |
| | total % cover | 0.00 | 0.95 | 0.351 | |
| **GPP** | **warming** | -0.19 | -3.04 | **0.004** | ** |
| | removal | -0.06 | -0.95 | 0.347 | |
| | warming : removal | 0.03 | 0.50 | 0.619 | |
| | **PC1** | 0.11 | 2.85 | **0.007** | ** |
| | PC2 | 0.10 | 1.59 | 0.121 | |
| | **total % cover** | 0.01 | 2.97 | **0.006** | ** |
| | soil moisture | 0.02 | 2.10 | 0.157 | |
| **ER** | **warming** | -0.34 | -3.81 | **0.000** | *** |
| | removal | -0.07 | -1.23 | 0.228 | |
| | warming : removal | -0.02 | -0.45 | 0.658 | |
| | **max air temp** | -0.05 | -2.14 | **0.039** | * |
| | **total % cover** | 0.00 | 2.39 | **0.022** | * |

**Table S3.** Linear mixed effect models summary table for the treatment effects (warming: W, removal: R) on C fluxes separately (NEE, ER, GPP) for the Hawkweed site. The A treatment corresponds to uninvaded ambient plots, while AC are ambient and invaded plots. The number of asterisks denote the significance: '***' when $p < 0.001$, '**' when $p < 0.01$ and '*' when $p < 0.05$.

| | Hawkweeds site - **Treatment effects** | | | | |
|---|---|---|---|---|---|
| **C flux** | **predictor** | $X^2$ | **Df** | ***p* value** | **significance** |
| NEE | intercept | 1.3 | 1 | 0.263 | |
| | treatment | 3.2 | 5 | 0.671 | |
| GPP | **intercept** | 98.1 | 1 | **<2e-16** | *** |
| | treatment | 7.1 | 5 | 0.215 | |
| ER | **intercept** | 96.9 | 1 | **<2e-16** | *** |
| | treatment | 5.2 | 5 | 0.389 | |

**Table S4.** Linear mixed effect models summary table for the treatments and predictors of C fluxes separately (NEE, ER, GPP) for the hawkweeds site. The number of asterisks denote the significance: '***' when $p < 0.001$, '**' when $p < 0.01$, '*' when $p < 0.05$ and '.' when $p < 0.1$. PC1 and PC2 were the two principal components extracted from the PCA analysis. PC1 summarizes the invaders biomass and traits, while PC2 has the most contribution from species diversity. $FD_Q$ and CWM RWC make reference to the calculated functional diversity and the community weighted means of relative water content (RWC).

| Hawkweeds site | | | | | |
|---|---|---|---|---|---|
| **C flux** | **Predictor** | **Estimate** | ***t* value** | **Pr(>\|*t*\|)** | **significance** |
| **NEE** | warming | 0.52 | 1.46 | 0.155 | |
| | removal | -0.09 | -0.48 | 0.637 | |
| | warming : removal | -0.02 | -0.11 | 0.917 | |
| | max air temp | 0.11 | 1.10 | 0.278 | |
| | PC1 | -0.25 | -1.86 | 0.071 | . |
| | **PC2** | -0.47 | -3.03 | **0.005** | ** |
| **GPP** | warming | -0.05 | -1.01 | 0.319 | |
| | removal | -0.07 | -1.31 | 0.200 | |
| | warming : removal | 0.01 | 0.16 | 0.876 | |
| | $FD_Q$ | -0.23 | -0.54 | 0.595 | |
| | **PC1** | 0.15 | 4.07 | **0.000** | *** |
| | **PC2** | 0.10 | 2.27 | **0.030** | * |
| **ER** | warming | 0.07 | 1.10 | 0.282 | |
| | **removal** | -0.09 | -2.54 | **0.016** | * |
| | warming : removal | 0.01 | 0.34 | 0.733 | |
| | max air temp | 0.02 | 1.11 | 0.276 | |
| | PC2 | -0.04 | -1.30 | 0.202 | |
| | **CWM RWC** | -0.02 | -2.82 | **0.008** | ** |
| | **evenness** | -1.05 | -2.48 | **0.018** | * |
